# Efficient and scalable Python implementation of ANCOM-BC for omics differential abundance testing

**DOI:** 10.64898/2026.01.26.701398

**Authors:** Zijun Wu, Huang Lin, James T. Morton, Qiyun Zhu

**Affiliations:** Biodesign Center for Fundamental and Applied Microbiomics, Arizona State University, 727 E. Tyler St., Tempe, AZ 85281, USA; Department of Epidemiology and Biostatistics, School of Public Health, University of Maryland, 48314 Paint Branch Dr, College Park, MD 20742, USA; Gutz Analytics, LLC, Boulder, CO, USA; School of Life Sciences, Arizona State University, 1151 S. Forest Ave., Tempe, AZ 85281, USA

**Keywords:** Differential abundance analysis, compositional data, microbiome, Python, benchmarking

## Abstract

ANCOM-BC (Analysis of Compositions of Microbiomes with Bias Correction) is a critical algorithm for differential abundance analysis. Designed for handling compositional data, it features the ability to estimate the unknown sampling fractions and correct the bias induced by their differences. While the original R package is robust for standard dataset sizes, the increasing magnitude of modern multi-omics studies presents computational challenges, often resulting in runtime bottleneck for high-dimensional data. Here, we present a native Python implementation of ANCOM-BC in the scikit-bio package that significantly accelerates the workflow, enabling its application to large-scale datasets. By leveraging the vectorization with NumPy and numerical optimization in SciPy, we enhanced the efficiency without compromising the statistical accuracy of the method. The benchmarking results revealed a dramatic increase in performance in both simulated and real datasets. We demonstrated that our Python implementation generates test results that match the original R package, while achieving a 100-fold acceleration in runtime and an 87% reduction in memory consumption. This new implementation allows for the analysis of datasets that are significantly larger than previously possible, and enables applications to alternative omics data types and complex statistical models that vary greatly by the number of samples, features, and covariates.

## Introduction

Differential abundance (DA) is an important problem for omics-wide association studies under different conditions due to the complexity of high-throughput data. Analysis of Compositions of Microbiomes with Bias Correction (ANCOM-BC) is a foundational and widely-used method for DA analysis in microbiome research (Lin and Peddada, 2020). A key strength of ANCOM-BC is its ability to explicitly estimate unknown sample-specific sampling fractions and to correct the bias introduced by their variation across samples. This provides a statistically grounded approach to addressing confounders from unobserved absolute abundances, a common challenge in microbiome studies, where measurements are typically limited to relative abundances. By addressing the source of bias across samples, ANCOM-BC enables more accurate inference of absolute feature abundances from compositional (i.e., relative abundance) data (Quinn *et al*., 2018). Similar limitations regarding compositional constraints apply to a variety of omics research fields, such as single-cell transcriptomics (Büttner *et al*., 2021), metabolomics (Krutkin *et al*., 2025), and glycomics (Bennett *et al*., 2025). Therefore, ANCOM-BC provides improved control of false discovery rates (FDRs) and greater interpretability compared with approaches that rely solely on relative abundances, making it an essential tool for rigorous and reliable DA analysis across the omics spectrum.

As omics workflows continue to scale in data complexity and integrate with machine learning paradigms, the impact of DA methods increasingly depends on their computational efficiency. In practice, DA serves not merely as a final statistical test, but as a repeated procedure of sensitivity assessment, cross-validation (Pelto *et al*., 2025), and dynamic feature selection step in predictive modeling (Romano *et al*., 2025) (Quinn and Erb, 2020). Consequently, efficient and scalable implementations are essential due to the increased computational cost in large-scale studies, enabling reproducible and flexible pipelines. This need motivates the development of Python-native implementations that ensure both computational enhancement and statistical accuracy.

Although bioinformatics software has traditionally been developed in R often, there has been a growing shift toward Python-based tools in recent years. Notable examples include the PyDESeq2 package (Muzellec *et al*., 2023) for differential gene expression (DGE, which is fundamentally equivalent to DA) analysis and the Scanpy framework for single cell analysis (Wolf *et al*., 2018). This transition is driven by several advantages of the Python language such as the access to well-maintained and efficient scientific computing libraries including NumPy and SciPy, the integration with machine learning and deep learning frameworks like scikit-learn and PyTorch, and the ability to reach a broader academic and industrial user base. Despite these benefits, no Python-native package currently exists for DA analysis with bias correction of microbiome data. A common workaround is to rely on Python-to-R interfaces, for example in the QIIME 2 platform (Bolyen *et al*., 2019), whereby R software is invoked from a Python environment. However, this approach complicates maintenance and deployment, and introduces computational overhead due to repeated data conversion and transfer between environments.

To address these limitations and fully leverage the advantages of Python-based software, we introduce a native Python implementation of the DA analysis, ANCOM-BC, originally proposed by Lin et. al (Lin and Peddada, 2020) and implemented in R. Using systematic benchmarking, we show that the Python implementation of ANCOM-BC delivers significantly enhanced computational performance while preserving statistical accuracy.

### Implementation

#### Algorithmic improvements

The key optimization in the Python implementation of ANCOM-BC focuses on efficiently solving the constrained optimization problem required to estimate the sampling bias inherent in compositional data. The algorithm assumes sparse effects on differential abundance and iteratively solves for the bias term that minimizes the discrepancy in the log-ratios of abundances. The acceleration of our Python implementation is achieved through a comprehensive re-engineering of this bias estimation algorithm, transitioning from iterative processing to a fully vectorized framework. We used efficient linear algebra backends in NumPy on the entire input matrix simultaneously with detailed analysis of the underlying mathematical derivations and the original implementation. In particular, we applied singular value decomposition (SVD) just once to replace iterative maximum likelihood estimation, and utilized Einstein summation extensively to avoid taxon-wise bias estimation. While eliminating the computational cost of iterations, our code retains numerical accuracy identical to the original method. The rigorous mathematical translation is supported by SciPy’s efficient numerical optimizer for the variance estimation of parameters of a Gaussian mixture model in the constrained bias-correction step, ensuring robust convergence without heavy computational redundancy. Additionally, the codebase is distinguished by its maintainability. We have intensive discussions and mathematical derivations within the in-line comments, designed to facilitate future development as well as AI readability. The implementation is further improved by comprehensive unit tests that validate statistical correctness and handle edge cases, allowing the tool to remain scientifically reliable.

#### Software features

An ‘ancombc’ function has been added to the Python package scikit-bio, with a simplified and versatile interface compared with the original R command ‘ANCOMBC’. Specifically, the Python function ‘ancombc’ consumes a “table-like” object that supports various common formats, such as BIOM tables (McDonald *et al*., 2012), AnnData objects (Wolf *et al*., 2018), Pandas and Polars dataframes, and NumPy arrays. The statistical model is defined by providing a metadata table as a dataframe, and an R-styled formula allowing for complex model definitions, such as the combination of categorical and numeric metadata columns and the inclusion of interaction terms. The behavior of the core ANCOM-BC algorithm is controlled by a small set of parameters, while the flexibility of data preprocessing is specified by the user. A comprehensive tutorial with sample data and code is included in the documentation of ‘ancombc’ to introduce the usage of this function together with data preprocessing and result postprocessing techniques to permit analysis consistent with the R package. Additionally, a function ‘struct_zero’ was added to test for structural zeros–features that are systematically absent from certain sample groups. This test was introduced along with the ANCOM-II method (Kaul *et al*., 2017) and has been incorporated into the R package ANCOMBC as part of the analytical workflow. In scikit-bio, we made it a standalone function as it is generally useful to the analysis of sparse omics data, with or without connecting to the ANCOM-BC test. These two functions were placed along with an existing set of DA testing methods, including ‘ancom’, the predecessor of ANCOM-BC (Mandal *et al*., 2015), and ‘dirmult_ttest’, the Python re-implementation of an underlying method of another widely adopted DA package: ALDEx2 (Gloor *et al*., 2025). They share consistently styled interfaces to permit user experimentation with different methods. Scikit-bio also provides a comprehensive suite of compositional data manipulation tools, such as log-ratio transformation and multiplicative replacement, to facilitate power users to build customizable DA analysis workflows.

### Performance comparison on synthetic data

We evaluated the computational performance of our Python code in comparison with the original ANCOM-BC R package using synthetic datasets generated by the built-in simulator of the R package. Specifically, we employed a Poisson lognormal (PLN) model that incorporates 10 continuous and 10 binary categorical covariates. This two-step simulation first sampled a latent Gaussian vector to establish taxon-to-taxon correlations and over-dispersion, then drew observed counts from a Poisson distribution conditioned on the exponentiated latent values and a scaling factor. The variance-covariance matrix was estimated from the Quantitative Microbiome Project (QMP) dataset (Vandeputte *et al*., 2017) with 106 samples and 91 OTUs. The simulator finally outputs a 10,000 samples by 10,000 features count matrix along with 20 covariates metadata. The performance assessment was stratified into three distinct scenarios to isolate the impact of sample size, feature dimensionality, and covariate complexity. First, we assessed sample complexity by fixing the feature count to 1,000 and covariates to 2, while incrementally increasing the number of samples. Second, we evaluated feature scalability by maintaining 1,000 samples and 2 covariates while varying the number of features. Finally, we examined the computational cost of model complexity by fixing the sample size to 1,000 and using a minimal feature set, while increasing the number of covariates. For each scenario, we recorded both the runtime and peak memory usage, providing a direct comparison of resource utilization between the two methods under increasing computational loads.

Based on the benchmarking results across all tested dimensions, the Python reimplementation of ANCOM-BC achieves orders-of-magnitude acceleration in runtime alongside substantial reduction in peak memory allocation compared to the original R package (**Figure 1**). The Python version scales exceptionally well with high-dimensional data, reaching a peak speedup of over 200× with 10,000 features (**Figure 1A**) and approximately 92× speedup of runtime with 10,000 samples (**Figure 1B**). This performance advantage extends to complex experimental designs, where the Python implementation maintains 30× to 46× speedup in covariate-heavy settings (**Figure 1C**). Complementing these runtime gains is a comparable improvement in memory efficiency. The Python implementation decreases peak memory consumption by a 7- to 30-fold reduction for large datasets (**Figure 1D, E**). Notably, more than 400-fold reduction is observed in smaller datasets with 10 samples and 1,000 features, possibly due to the demand of initialization in R (**Figure 1E**). Collectively, the significant decreases of both runtime and memory usage in Python implementation of ANCOM-BC validate its efficiency in analyzing massive omics datasets on standard hardware.

**Figure 1.**
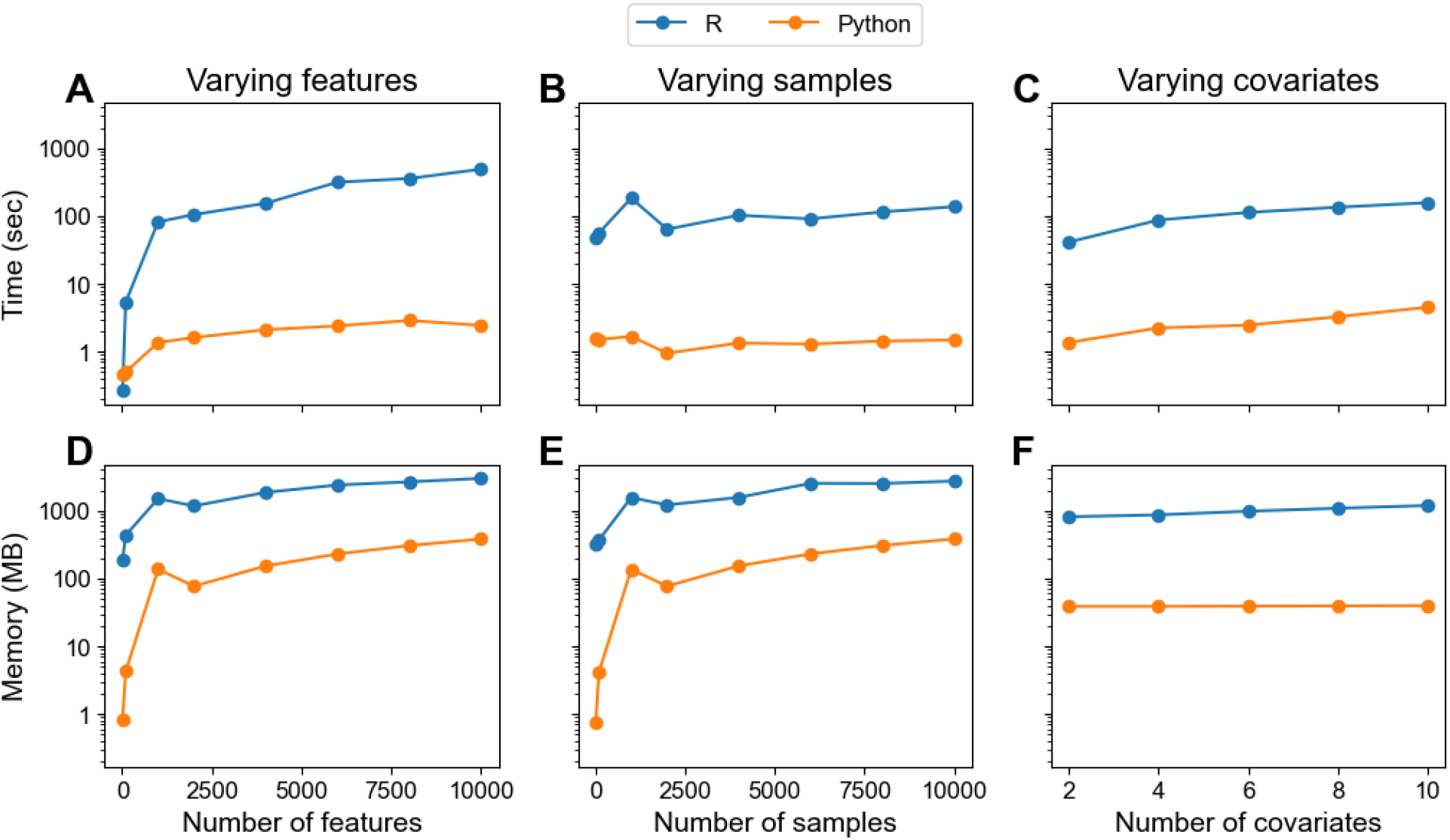
Benchmark results averaged over three repeated runs, using a single thread for each package. Runtime of R and Python versions of ANCOM-BC using synthetic dataset varying the number of features **(A)**, samples **(B)**, and covariates **(C).** Memory usage comparison between R and Python version of ANCOM-BC on synthetic dataset with different number of features **(D)**, samples **(E)**, and covariates **(F)**.

### Applications on real-world omics datasets

To evaluate the Python implementation of ANCOM-BC across diverse biological contexts and data dimensions, we utilized three distinct datasets representing varying degrees of sparsity and complexity. The HITChip Atlas dataset (Lahti *et al*., 2014), demonstrated in the official ANCOM-BC tutorial, includes 873 samples and 21 features modeled against 7 covariates after preprocessing. To assess the performance in high-dimensional microbiome settings, we employed a filtered shotgun metagenomic dataset from the Earth Microbiome Project 500 (EMP500) (Shaffer *et al*., 2022) comprising 753 samples and an expanded feature space of 8,351 taxa, adjusted for 2 covariates. Finally, extending the framework beyond microbiomics, we included a single-cell RNA-seq subset from an atherosclerosis study that contains 7,826 cells, 1,226 genes, and a complex experimental design involving 24 covariates. As shown from Figure 2, the Python implementation returns almost identical results of differential abundant features (**Figure 2A, B, C**) while achieving greatly accelerated speed and reduced memory usage (**Figure 2D, E**).

**Figure 2.**
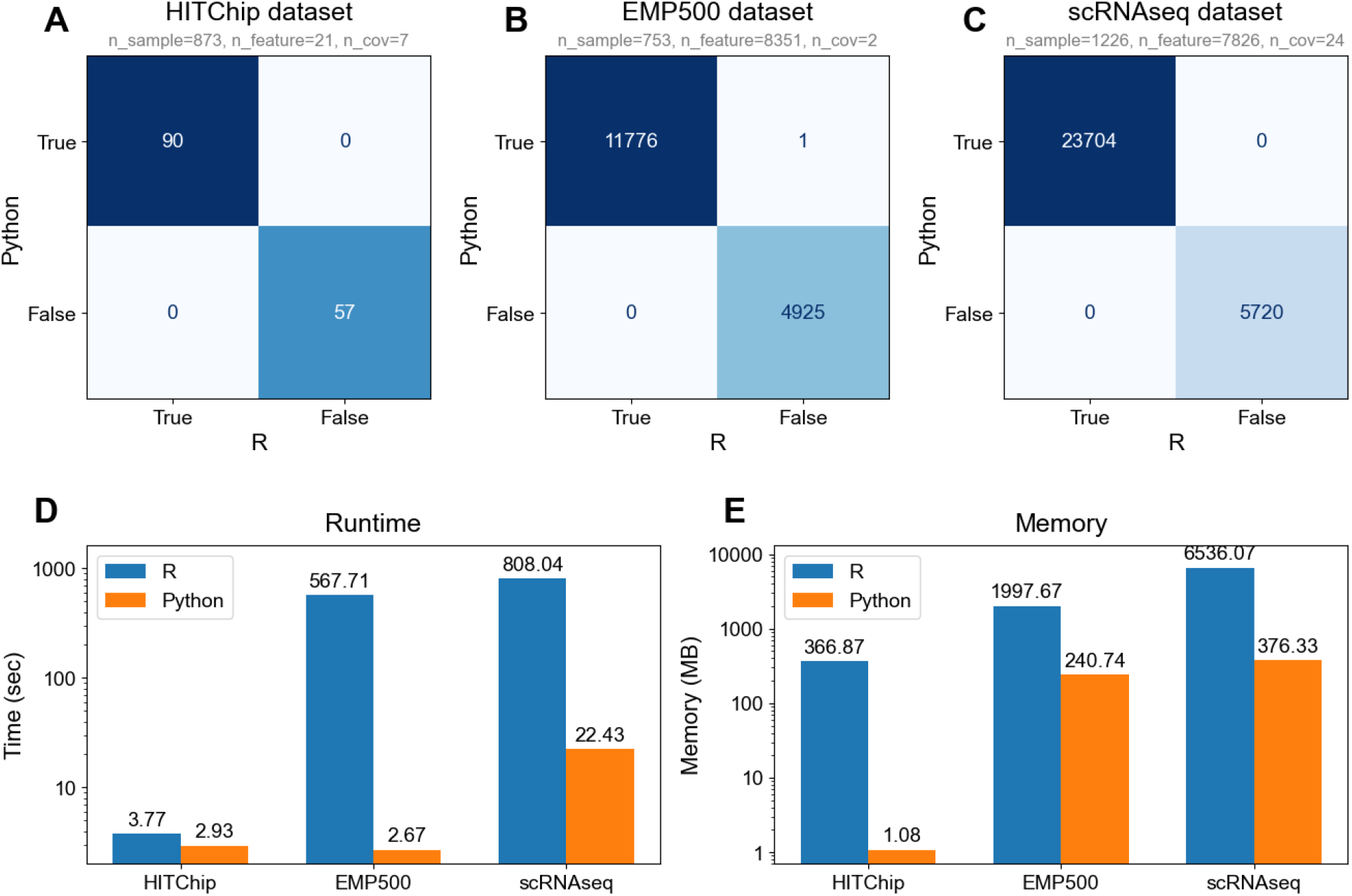
Differential abundance test results according to ANCOM-BC and Python implementation on **(A)** HITChip Atlas dataset, **(B)** EMP500 dataset and **(C)** single cell RNAseq dataset on atherogenesis study. (**D**) Runtime and (**E**) memory usage comparison.

## Conclusion

The Python implementation of the ANCOM-BC algorithm in scikit-bio provides a robust and computationally efficient framework for differential abundance (DA) analysis. By introducing this new feature, we aim to bridge a critical gap in the Python omics ecosystem and facilitate the integration of modern data science workflows into microbiome studies and beyond. With the enhanced performance of ANCOM-BC, we unlock its potential as an explainable AI technique, particularly through the identification of discriminative balances (Quinn and Erb, 2020). Coupling differential abundance to machine learning models imposes a massive computational burden, high-efficiency implementations are therefore essential for accessing feature importance by sensitivity analysis and cross-validation. The future landscape of Python-based DA analysis is set to major improvements driven by the needs for complex statistical modeling and high-performance computing. Concurrently, we are working on advanced statistical modeling including variance regularization, sensitivity analysis, and support for multigroup analysis (Lin and Peddada, 2024). Upcoming developments will also prioritize sparse matrix operations for increasingly massive multi-omics datasets, as well as data structure compatibility for an end-to-end bioinformatics workflow.

## Code availability

The Python implementation of ANCOM-BC has been incorporated into the scikit-bio package, available at: https://scikit.bio. The source code of scikit-bio is licensed under BSD-3 and hosted at the public GitHub repository: https://github.com/scikit-bio/scikit-bio. Version 0.7.1 of scikit-bio, which started to ship the ANCOM-BC algorithm, has been permanently archived at the Zenodo repository: https://zenodo.org/records/17487690. The benchmark datasets and scripts are available at: https://github.com/iiiime/ancom-bc_bm.

## Author contributions statement

Z.W. and Q.Z. conceived the study and implemented the software code. Z.W. performed testing and benchmarking, analyzed the results, and led the writing of the manuscript. H.L. and J.T.M. provided advice on the scope and design of the algorithm. All authors contributed to the writing and review of the manuscript.

## Acknowledgements

This work is sponsored by the U.S. Department of Energy, Office of Science under award DE-SC0024320.

